# Peripheral microtubules ensure asymmetric furrow positioning in neural stem cells

**DOI:** 10.1101/2020.09.10.291112

**Authors:** Alexandre Thomas, Emmanuel Gallaud, Aude Pascal, Laurence Serre, Isabelle Arnal, Laurent Richard-Parpaillon, Matthew Scott Savoian, Régis Giet

## Abstract

Neuroblast (NB) cell division is characterized by a basal positioning of the cleavage furrow resulting in a large difference in size between the future daughter cells. In animal cells, furrow placement and assembly is governed by centralspindlin, a highly conserved complex that accumulates at the equatorial cell cortex of the future cleavage site and at the spindle midzone. In contrast to model systems studied so far, these two centralspindlin populations are spatially and temporally separated in NBs. A cortical leading pool is located at the basal cleavage furrow site and a second pool accumulates at the midzone before travelling to the site of the basal cleavage furrow during cytokinesis completion. By manipulating microtubule (MT) dynamics, we show that the cortical centralspindlin population requires peripheral astral microtubules and the Chromosome Passenger Complex (CPC) for efficient recruitment. Loss of this pool does not prevent cytokinesis but enhances centralspindlin levels at the midzone leading to furrow repositioning towards the equator and decreased size asymmetry between daughter cells. Together these data reveal that the asymmetrical furrow placement characteristic of NBs results from a competition between spatially and functionally separate centralspindlin pools in which the cortical pool is dominant and requires peripheral astral microtubules.

## Introduction

Cytokinesis in somatic cells ensures the equal partitioning of the segregated chromosomes and is responsible for the division of the mother cell’s cytoplasm into two daughters. This process requires the highly orchestrated assembly and constriction of an acto-myosin contractile ring, usually at the cell’s center. The use of various model systems has clearly established that the mitotic spindle defines the position of the contractile ring and the resulting cleavage furrow^1, 2^. Two populations of mitotic spindle microtubules (MTs) have been shown to trigger the assembly of the contractile machinery during late anaphase. The first is a sub-population of astral MTs. These MTs emanate from the centrosomes to the equatorial cortex where they deliver furrow-inducing signals ^3-5^. The second population, comprises the spindle midzone, a region of antiparallel MT overlap and interdigitation within the central spindle that assembles between the decondensing daughter nuclei. In many symmetrically dividing somatic cell types the relative contribution of these two populations has been difficult to unambiguously determine due to their close proximity at the cell’s equator. Yet, experiments during the last few decades have helped to propose a common mechanism across model systems in which the furrow-inducing signals emanate from both cortical proximal astral MTs and the spindle midzone with each acting in parallel. However, these pathways do not appear to be equivalent. For instance, if the furrow is located away from the midzone, it will regress and a new one will be established at the midzone location ^1, 3, 6, 7^. Thus, in equatorially dividing cells, the spindle midzone pathway acts dominantly and can reset furrowing.

Centralspindlin is the main orchestrator of furrowing. This protein complex is a tetramer composed of two subunits of the Kinesin 6 (Pavarotti-klp in *Drosophila melanogaster*) and two subunits of the MgcRacGAP (Tumbleweed in *Drosophila melanogaster*). Tumbleweed is essential for the activation of the Rho-GEF Ect2 (Pebble in *Drosophila melanogaster*). The formation of Rho-GTP triggers the local activation of Rho Kinase and phosphorylation of non-muscle Myosin Regulatory Light Chain, an event that stimulates myosin activation and ultimately drives cytoplasmic cleavage ^8, 9^. While it is well established that centralspindlin acts along central spindle MTs and accumulates at the cell cortex equator to promote symmetrical cleavage, far less is known about how this complex governs asymmetrical divisions. *Drosophila* neural stem cells (Neuroblasts, NBs) are characterized by a biased furrow placement towards the basal region of the cell. Asymmetric cytokinesis triggers the formation of a large apically positioned cell that retains the NB identity, and a small basal ganglion mother cell (GMC) that will undergo differentiation ^10^. Asymmetry is apparent prior to furrow initiation and can be detected during early anaphase as myosin redistributes from around the cortex to a more basal position ^11^. Previous studies have shown that this process is under the strict control of the NB polarity machinery but is also influenced by the spindle midzone and the Chromosome Passenger Complex (CPC) ^12, 13^. To better understand the mechanism of furrow positioning in asymmetrical cytokinesis, we have genetically manipulated spindle size and MTs dynamics in *Drosophila* NBs. Our data indicate that the mechanisms dictating asymmetrical daughter cell size are extremely robust and tolerate increases in spindle length and shape. We report that furrowing initiates away from the midzone, in a basal position through the action of a subcortical centralspindlin pool targeted by peripheral astral microtubules. When these MTs are ablated, centralspindlin recruitment at the furrow is impaired and becomes abnormally enriched at the midzone causing repositioning of the cleavage site, thus affecting the size asymmetry of the daughter cells. Together these results reveal that unlike most systems, in *Drosophila* NBs that are characterized by a high level of cell size asymmetry during cell division, a population of peripheral astral MTs, and not the spindle midzone, defines and maintains asymmetric cleavage furrow positioning.

## Material and methods

### Molecular biology

Msps cDNA was provided by G. Rogers (University of Arizona, USA), amplified by PCR and inserted into pENTR (Life Technologies) to generate the pENTR-Msps entry clone. pENTR-Ensc has been previously described ^14^. The pENTR-Feo and the pENTR-Tum entry clones were obtained from P.P D’Avino (University of Cambridge, UK). The pENTR-Feo and pENTR-Msps entry clones were each subsequently recombined into pTWV and pTWR (Carnegie Institute, USA) to generate constructs allowing the expression of Feo-VenusFP and Msps-RFP fusion proteins, respectively, under the control of the GAL4 protein using the Gateway recombination cloning technology (Life Technologies). pENTR-Ensc was recombined into pDEST-MBP (a gift from H. Ohkura, University of Edinburgh, UK) to allow the expression of a recombinant Ensconsin protein with a C-terminal MBP tag.

### Fly stocks

All flies were maintained under standard conditions at 25°C. The *ensconsin* mutant fly stocks *ensc*Δ*null* and *ensc*Δ*N*, referred to *ensc* flies, were characterized previously (Sung et al 2008). UAS-Ensconsin-Venus transgenic flies, overexpressing Ensconsin-Venus, have been described ^15^. UAS-Feo-Venus and UAS-Msps-RFP overexpressing flies were obtained from BestGene (USA) following P-element mediated transformation. UAS-Klp10A flies were supplied by C. Dahmann (Max Planck Institute, Germany) ^16^. UAS-Klp67A (ID # F001232) stock was obtained from FlyORF ^17^. UAS-Klp67A-RNAi (VDRC ID 52105) and UAS-Mad2-RNAi (VDRC ID 106003) transgenic fly lines were obtained from the Vienna Drosophila RNAi Center ^18^. Sqh-GFP ^19^, UAS-GFP-Pav-klp and Ubiquitin-βtub-GFP expressing flies ^20, 21^ were supplied by R. Karess (Institut Jacques Monod, France) and by D. Glover (University of Cambridge, UK), respectively. The Pavarotti mutant *pav*^*B200*^ flies were obtained from E. Montembault (Institut Europeen de Chimie et Biologie, France) ^22^ and Survivin mutant allele *svn*^*2180*^ flies were courtesy of Jean-René Hyunh (College de France, France) ^23^. Flies expressing the membrane-localized PH-PLCδ-GFP and PH-PLCδ-RFP proteins were provided by A. Guichet (Institut Jacques Monod, France) ^24, 25^. RFP-Tubulin flies were provided by R. Basto (Institut Currie, France). The GFP-AurA expressing fly stock was described previously ^26^. The following stocks were obtained from the Bloomington Stock Center: Feo-GFP expressed under the ubiquitin promoter (BDSC 59273, ^27^), *sas-4*^*s2214*^ mutant (BDSC 12119, ^28^), 69B-Gal4 (BDSC 1774), Insc-Gal4 (BDSC 8751), UAS-mCherry-α-tubulin (BDSC 25774 and BDSC 25773). The 69B-Gal4 fly stock was used to drive over-expression in the fly CNS for the following UAS regulated transgenes: Ensconsin, Klp67A, Msps, GFP-Pav-Klp together with *Mad2* RNAi and UAS-mCherry. The Insc-Gal4 strain was used to drive over-expression of Ensconsin, Klp10A, UAS-GFP-Pav-klp, Feo-Venus and mCherry-α tubulin transgenes in the central brain.

### Production of recombinant proteins

MBP and Ensconsin-MBP were induced in *E.coli*, for 4 h at 25°C. The proteins were purified on amylose column as described by the manufacturer (BioLabs) and stored in small aliquots at −80°C.

### TIRF microscopy and analysis of MT dynamics

Tubulin was purified from bovine brain and fluorescently labeled with ATTO 488 and ATTO 565 or biotinylated as described before ^29, 30^. Briefly, microtubule seeds were prepared from biotinylated and ATTO-565-labeled tubulin in the presence of Guanosine-5’-[(α,β)-methyleno]triphosphate (GMPCPP) in BRB80 buffer (80 mM Pipes, 1 mM EGTA, 1 mM MgCl_2_, pH 6.74) ^30^. Flow chambers were prepared with functionalized silane-PEG-biotin coverslips and silane-PEG glass slides, as previously described (Ramirez-Rios et al., 2017). The chamber was successively perfused at room temperature with neutravidin (25 µg/ml in 1% BSA in BRB80), PLL-g-PEG (2 kD, 0.1 mg/ml in 10 mM Hepes, pH 7.4), BSA (1% in BRB80 buffer) and microtubule seeds. The following assembly mixture was then injected: 14 µM tubulin (containing 15 % ATTO-488-labeled tubulin) without or with 200 nM MBP or MBP-Ensconsin in TIRF assay buffer (4 nM DTT, 50 mM KCl, 1% BSA, 1 mg/mL glucose, 70 µg/mL catalase, 580 µg/mL glucose oxidase, 0.05% methylcellulose (4000 centipoise) in BRB80). Time-lapse images were recorded at 35°C at a rate of one frame per 5 seconds on an inverted Eclipse Ti Nikon microscope equipped with an Apochromat 60×1.49 N.A oil immersion objective, an iLas^2^ TIRF system (Roper Scientific), and a cooled charge-coupled device camera (EMCCD Evolve 512, Photometrics) controlled by MetaMorph 7.7.5 software. Microtubule dynamic parameters were analyzed in Image J on kymographs obtained using an in-house KymoTool macro (available upon request to). Growth and shrinkage rates were determined from the slopes of microtubule growth and shrinkage phases. The catastrophe and rescue frequencies were calculated by dividing the number of events per microtubule by the time spent in growing and shrinking states, respectively.

### Antibodies and Western blotting

The following antibodies and concentrations were used in this study: polyclonal rabbit anti-Msps (1:5000) provided by J. Raff ^31^, polyclonal rabbit anti-Klp67A (1:500) supplied by G. Goshima ^32^, polyclonal rabbit anti-Klp10A (1:1000) was courtesy of G. Rogers ^33^ and rabbit anti-Myosin (1:2000) was provided by R. Karess ^34^. The anti-Ensconsin antibody raised against the Kinesin binding domain has been previously described^14^. Rabbit anti-PKCζ (C-20, 1:200) and anti-actin polyclonal antibodies (sc-1616, 1:5000) were obtained from Santa Cruz Technology. Monoclonal mouse anti-alpha Tubulin (clone DM1A, T2199; 1:500) and rabbit polyclonal anti-phosphorylated histone H3 (Ser10) (06570, 1:500) antibodies were obtained from Millipore. Monoclonal rat anti-Miranda antibody (ab197788, 1:1000) was obtained from Abcam. Secondary antibodies were labelled with either Alexa Fluor-conjugated (1:1000) or peroxidase-conjugated secondary antibodies (1:5000), each obtained from Life Technologies. For Western Blotting ECL reagents were purchased from ThermoFisher.

### Live cell microscopy

Third-instar larval brains were dissected in Schneider’s Drosophila medium supplemented with 10% FCS. Isolated brains were loaded and mounted on stainless steel slides, and the preparations were sealed with mineral oil (Sigma-Aldrich) as previously described ^14^. For MT depolymerization experiments, larval brains were incubated during 30 min in the above medium supplemented with colchicine at a final concentration of 15 μM. After incubation, brains were mounted and processed for live cell imaging.

Images were acquired at 25°C using a CSU-X1 spinning-disk system mounted on an inverted microscope (Elipse Ti; Nikon) equipped with a 60X 1.4 NA objective. At 20, 30 or 60 sec intervals 10 z-steps were acquired with 1µm intervals. Fluorescent protein probes were excited with 488nm or 561nm laser light and the images were captured using a sCMOS ORCA-Flash4.0 (Hamamatsu) camera. Recordings were controlled using MetaMorph acquisition software. Data were processed in ImageJ and viewed as maximum-intensity projections prior to analysis or figure preparation.

### Photo-ablation experiments

Photo-ablations were performed with a Mai-Tai two-photon infrared laser (Spectra Physics) attached to a Leica SP5 confocal microscope equipped with a 60X 1.3 NA objective with the stage maintained at 25°C. *Z*-series consisting of 10, 1µm steps were acquired before and after the photo-ablation at 30 sec intervals. Photo-ablation was performed on the basal centrosome using flies expressing GFP-H2A, PH-PLCδ-GFP and Aurora A-GFP. Anaphase onset was identified as the first signs of sister chromatid separation. The photo-ablation was considered complete after the Aurora A signal on the basal centrosome could no longer be detected.

### Live cell imaging analysis

Measurements of fluorescence intensities, mitotic spindle lengths and diameters of NB and GMC cells were performed with ImageJ software ^35^. The Sqh-GFP analyses were done on the maximum projection of two optical sections (1 μm). The polarity-dependent apical clearing was calculated, as the first time point after anaphase onset, when myosin began to disappear from the apical cortex ^36^. NB cortex curvature analyses were performed according to previously defined methods ^13^. The furrow shift was determined as the distance between the first ingression site and final cleavage site. To quantify the Myosin-GFP furrows width; a segmented line was drawn along the NB half-cell cortex during anaphase and the GFP intensity profiles were quantified along this line using ImageJ. The furrow width was measured as the relative half-cell cortex length containing 60% of the maximum Sqh-GFP signal intensity.

### Immunofluorescence analysis

Larval brains from each genotype were processed for immunofluorescence studies as described previously ^14^. Briefly, wandering third instar larval brains from were dissected in testis buffer (TB: 183 mM KCL, 47 mM NaCl, 10 MM Tris, and 1 mM EDTA, pH 6.8) and brains were fixed for 20 minutes at 25°C in TBF (TB supplemented with 10% formaldehyde, and 0.01% Triton X-100). Brains were then washed twice in PBS for 15 minutes, and twice in PBS Triton X-100 0.1% for 15 minutes. The brains were first incubated for 60 minutes at 25°C in PBSTB (1% BSA), before incubation with secondary antibodies. The samples were observed with a SP5 confocal microscope (Leica) equipped with a 63X 1.4 NA objective lens. Images are maximum intensity projections consisting of 4 optical sections acquired at 0.5 µm intervals.

### Quantification of peripheral MTs in fixed NBs during mid anaphase

Z-series were acquired every 0.2 μm using a LSM 880 confocal microscope with Airyscan (Zeiss) for telophase NBs. Images were then processed with the Zen software. Images were analyzed with ImageJ as maximum intensity projections (0.8 μm) consisting of 5 optical (0.2 μm) sections in the plane of the furrow.

## Statistical analysis

Differences between datasets were assessed with Prism 7.0a software (GraphPad), either by non-parametric tests (Mann-Whitney-Wilcoxon) or parametric tests (Unpaired T). Non-significance (ns) threshold was when *P*>0.05.

## Results

### Cell size asymmetry is compromised following Ensconsin depletion but not over-expression in NBs

Neuroblasts divide asymmetrically to generate a large self-renewing neuroblast (NB) and a smaller differentiating ganglion mother cell (GMC, Figure 1 A. We previously showed that Ensconsin is required for MT polymerization during cell division; consequently *ensc* mutant spindles are shorter than their wild type (WT) counterparts ^14^. To investigate the possible consequences of a change in spindle length on NB asymmetric cell division, we first analyzed, by live cell imaging, cell size asymmetry of dividing NBs in wild type and *ensc* mutants (Figure 1). We confirmed the previous finding that loss of Ensconsin triggered a ∼10% decrease in mitotic spindle length (Figure 1 B, C). Strikingly, the *ensc* mutants displayed a small yet statistically significant reduction in the ratio between NB and GMC diameters indicating a loss of asymmetry (Figure 1 D). This defect could either result from the associated change in spindle length or indicate some uncharacterized function for Ensconsin in asymmetrical size fate determination. To further explore the role of Ensconsin in MT dynamics *in vitro*, we used TIRF microscopy and recombinant Ensconsin protein (Figure 1 E). Ensconsin-MBP had a small but significant effect on MT growth rate. Most striking was the ∼50% reduction in the rate of MT shrinkage and the more than 3 times increase in the rescue frequency compared to controls or MBP alone (Figure 1 F). In line with these results, over-expression of Ensconsin (Ensc-OE) in NBs lead to elongated spindles that buckled when reaching the cortex (Figure 1 G, H, S1), consistent with previous work in symmetrically dividing S2 cells ^14^. Despite the increase in MT polymerization and spindle length, the level of size asymmetry remained unperturbed following cytokinesis in Ensc-OE NBs (Figure 1 I).

**Figure 1.**
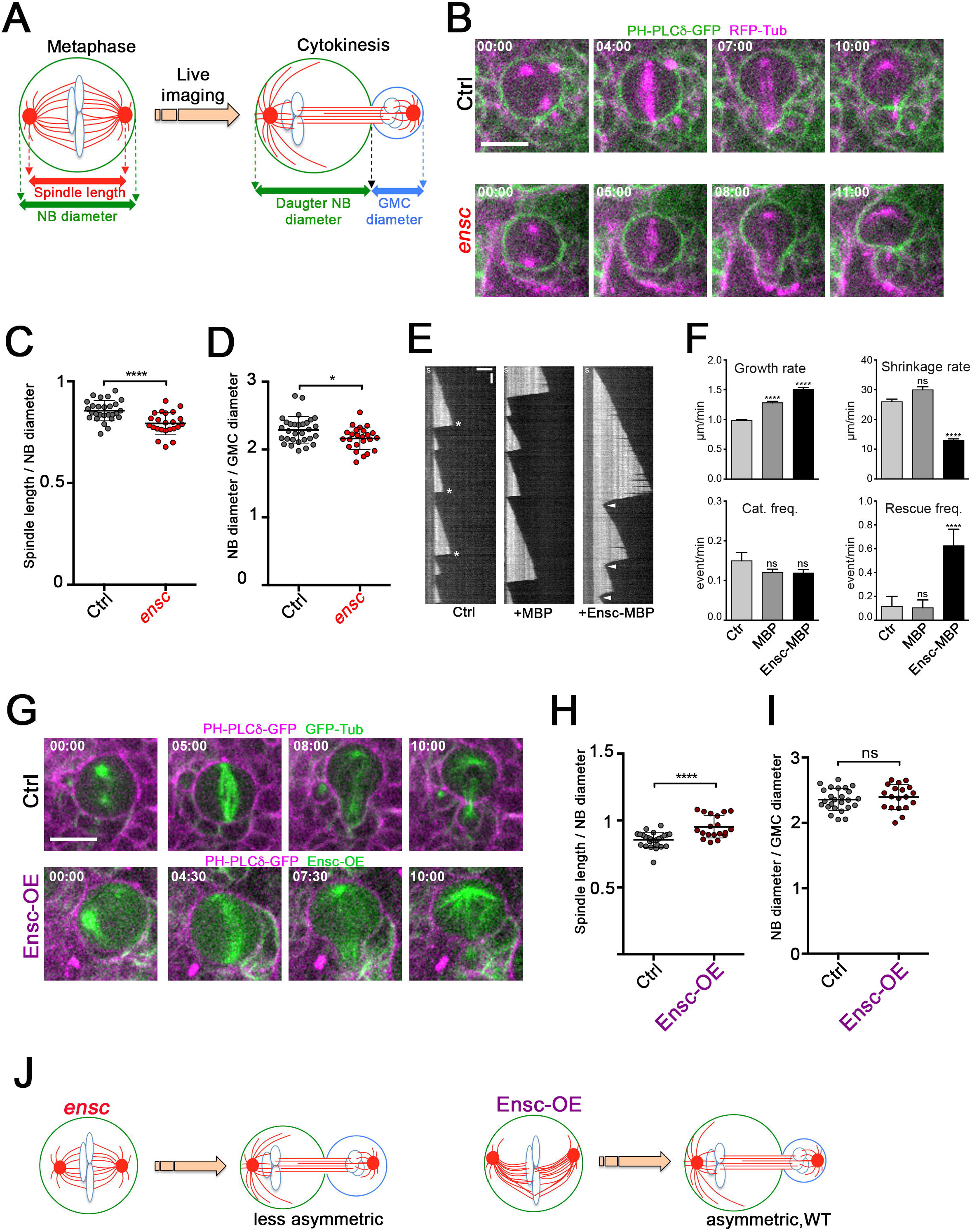
Analysis of cell size asymmetry in *ensc* and Ensc-OE NBs. A) Scheme of a NB during cell division during metaphase (left) and telophase (right). Note that the NB cell division is asymmetric and produces a large NB and a small Ganglion Mother Cell (GMC) during cytokinesis. B) Selected images of a control NB (top) and an *ensc* mutant NB (bottom) during cell division. The membranes are displayed in green and the MTs are displayed in magenta. Time is min:s. Scale bar: 10μm. C) Dot plot showing the mitotic spindle length/NB diameter ratio (± s.d.) in control (0.86±0.05, *n=*27) or in *ensc* NBs (0.79±0.06, *n=*23), ***: *P*<0.0001 (Wilcoxon test). D) Dot plot showing the NB diameter/GMC diameter ratio (± s.d.) in control (2.29±0.20, *n=*27) or in *ensc* NBs (2.16±0.20, *n=*23), *: *P*<0.05 (Wilcoxon test). E) Kymographs showing microtubules assembled from GMPCPP seeds and 14 µM tubulin in absence or in presence of 200 nM of MBP or MBP-Ensconsin. Horizontal and vertical scale bar are 5μm and 60s respectively. F) Graphs showing the growth and shrinkage rates and the catastrophe and rescue frequencies determined from kymographs shown in E. ns: non significant; ****: *P*<0.0001 (Kruskal-Wallis ANOVA followed by post-hoc Dunn’s multiple comparison, total number of growth events = 116, 123 and 107, shrinkage events = 67, 73 and 81, catastrophe events = 94, 96, 83 and rescue events = 3, 3 and 33 for the control, MPB and MPB-ensconsin respectively). G) Selected images of a control NB (top) and an Ensc-OE NB (bottom) during cell division. Membranes are displayed in magenta and MTs or Ensconsin are displayed in green. H) Dot plot showing the mitotic spindle length/NB diameter ratio (± s.d.) in control (0.86±0.06, *n=*25) or in Ensc-OE NBs (0.95±0.08, *n=*19), ****: *P*<0.0001 (Wilcoxon test). I) Dot plot showing the NB diameter/GMC diameter ratio (± s.d.) in control (2.36±0.17, *n=*25) or in Ensc-OE NBs (2.40±0.19, *n=*19), ns: non-significant (Wilcoxon test) J) Summary of NB division in *ensc* (left) or Ensc-OE (right). *ensc* mutant NBs display shorter spindles and undergo less asymmetric cell division while Ensc-OE NBs, despite harboring long spindles, divides asymmetrically similar to WT. Time is min:s.

### Enhancement of spindle length through over-expression of Msps or depletion of Kinesin-8 MT depolymerase does not alter cell size asymmetry

To determine if daughter cell size asymmetry is insensitive to stimulation of MT growth, we quantified size asymmetry following over-expression of the microtubule associated protein Mini spindles, the fly orthologue of MAP215/ch-TOG, a protein with MT polymerization properties ^37-39^. In parallel, we performed RNAi-mediated depletion of the MT depolymerizing Kinesin-8 fly family member Klp67A. This kinesin depolymerizes microtubules and its depletion leads to the formation of exceptionally long spindles in *Drosophila* cells ^40, 41^. Similar to Ensc-OE, over-expression of Msps-RFP (Msps-OE) or RNAi-mediated depletion of Klp67A led to the formation of long and bent mitotic spindles (Figure 2 A, B and D; Figure S1; Video 1 and 2). Neither perturbation affected the post-cleavage asymmetrical cell size (Figure 2A, C and E). These data suggest that asymmetric cell size regulation is not sensitive to an increase in MT polymer or spindle length elongation.

**Figure 2.**
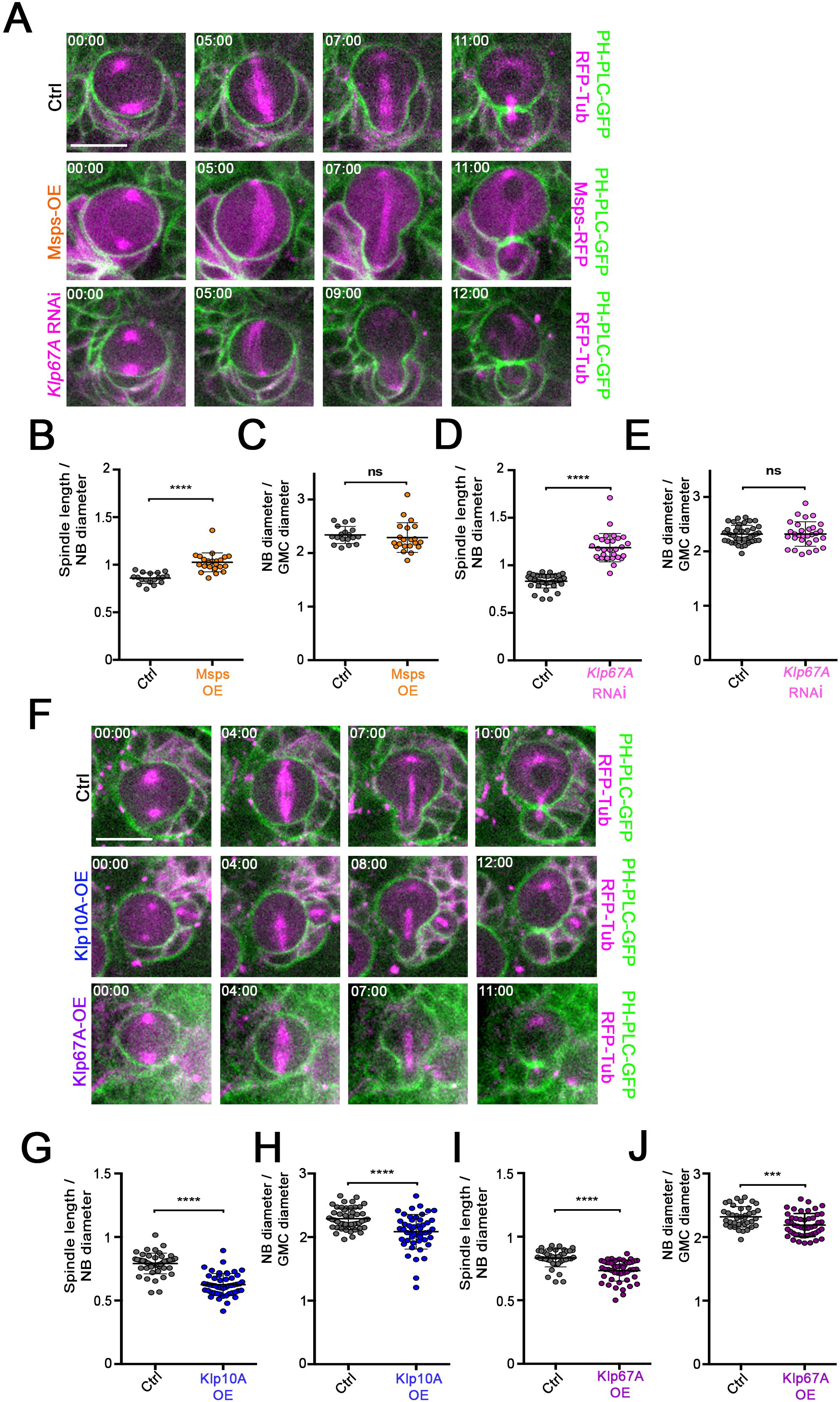
Analysis of cell size asymmetry in NB following modification of several MAP protein levels. A) Selected images of control (top), Msps-OE (middle) and *Klp67A* RNAi (bottom) NBs. The membranes are shown in green, the MTs (top and bottom) and Msps (middle) are shown in magenta. Time is min:s. Scale bar: 10μm. B) Dot plot showing the mitotic spindle length/NB diameter ratio (± s.d.) in the NB of control (0.86±0.06, *n=*18) or in Msps*-*OE transgenic flies (1.03±0.1, *n=*23), ****: *P*<0.0001 (Wilcoxon test). C) Dot plot showing the NB diameter/GMC diameter ratio (± s.d.) in control (2.34±0.16, *n=*18) or in Msps-OE NBs (2.29±0.28, *n=*21), ns: non-significant (Wilcoxon test). D) Dot plot showing the mitotic spindle length/NB diameter ratio (± s.d.) in control NBs (0.83±0.07, *n=*40) or in *Klp67A* RNAi NBs (1.19±0.15, *n=*30), ****: *P*<0.0001 (Wilcoxon test). E) Dot plot showing the NB diameter/GMC diameter ratio in control (2.31±0.16, *n=*40) or in *Klp67A* RNAi NBs (2.32±0.23, *n=*30), ns: non-significant (Wilcoxon test). F) Selected images of control (top), Klp10A-OE (middle) and Klp67A-OE (bottom) NBs. The membranes are shown in green, the MTs are shown in magenta. Scale bar: 10μm. Time is min:s. G) Dot plot showing the mitotic spindle length/NB diameter ratio (± s.d.) in control NBs (0.79±0.08, *n=*49) or in Klp10A-OE NBs (0.63±0.09, *n=*49), ****: *P*<0.0001 (Wilcoxon test). H) Dot plot showing the NB diameter/GMC diameter ratio (± s.d.) in control NBs (2.29±0.17, *n=*49) or in Klp10A-OE NBs (2.00±0.27, *n=*49), ****: *P*<0.0001 (Wilcoxon test). I) Dot plot showing the mitotic spindle length/NB diameter in control NBs (0.83±0.07, *n=*40) or in Klp67A-OE NBs (0.73±0.08, *n=*48), ****: *P*<0.0001 (Wilcoxon test). J) Dot plot showing the NB diameter/GMC diameter ratio in control NBs (2.32±0.16, *n=*40) or in *Klp67A* RNAi NBs (2.19±0.18, *n=*48), ****: *P*<0.0001 (Wilcoxon test).

### Spindle shortening through over-expression of Kinesin-8 or -13 family MT depolymerases decreases cell size asymmetry

To investigate if the size asymmetry reduction observed in *ensc* mutants (Figure 1) was unique to Ensconsin or rather a common effect of spindle shortening, we induced other perturbations of MT-polymerization by over-expressing two MT depolymerizing kinesins; either Klp10A which belongs to the Kinesin-13 family (Klp10A-OE) or the Kinesin-8 member Klp67A (Klp67A-OE) (Figure S1). Importantly, whereas the depletion of either causes spindle elongation, their over-expression results in abnormal shortening ^40, 42-45^. As predicted both Klp10A-OE and Klp67A-OE NBs exhibited shorter spindles although the length reduction was more pronounced in Klp10A-OE cells (Figure 2 F, G, I Video 3). Depending on the spindle’s length, it assumed a lesser or greater displacement relative to the cell center. Interestingly, in these shortened spindle cells, like with *ensc* mutants, we found that the NB/GMC diameter ratio was significantly impaired indicating that cell division was more symmetric compared to controls (Figure 2 H and J). Thus, defective MT polymerization leading to spindle shortening due to loss of Ensconsin function or over-expression of the Kinesin-8s and 13s biases asymmetric cell division.

### Defective MT growth leads to an apical shift of the basal cleavage furrow after anaphase onset

In asymmetrically dividing NBs, it was described that both polarity and midzone-dependent mechanisms ensure that furrow components, including myosin, are positioned basally to generate daughter cells of different sizes ^11, 13, 36^. To investigate the behavior of the furrow, we monitored the dynamics of the regulatory light chain of myosin in live NBs using Sqh-GFP ^46^. In control cells, we confirmed that Myosin-GFP was uniformly present at the cell cortex before anaphase (Figure 3 A and Video 4). In *ensc*, Klp67A-OE and Klp10A-OE NBs, the polarity-dependent step of myosin redistribution from the apical cortex was similar to controls, in agreement with the perturbations not altering cell polarization (Figure S2 A and B). We also analyzed the furrow positioning through curvature measurements of the cell membrane (^13^ and Figure 3 B). While the furrow position remained stably placed from anaphase until cytokinesis in control NBs, we found it shifted significantly towards the apical side during completion of cell division in *ensc* and Klp67A-OE cells, with a maximal displacement observed for Klp10A-OE NBs (Figure 3, C Video 5 and 6). Moreover, while the furrow width was consistently ∼10% of the half-cell cortex length in control NBs as revealed by Myosin-GFP (Figure 3 D and E), the signal occupied a larger space in *ensc* and Klp67A-OE NBs with the maximum width of ∼25% of the half-cell cortex length observed for Klp10A-OE cells at comparable time points (Figure 3 A, E, Video 6). From this we conclude that proper MT growth is required for maintaining furrow size and position during asymmetrical cell division.

**Figure 3.**
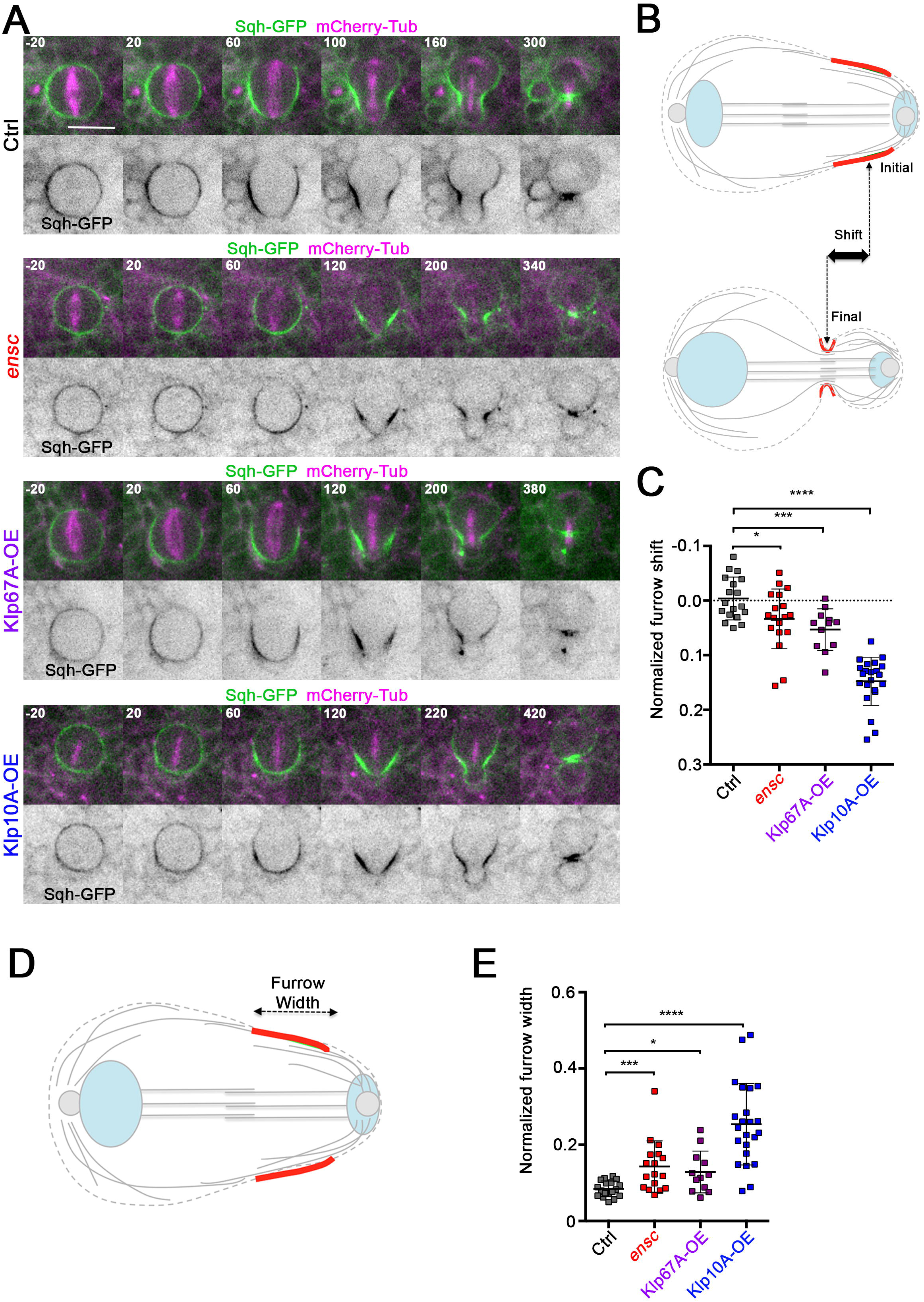
Analysis of myosin dynamics and furrow positioning in *ensc*, Klp67-OE and Klp10A-OE. A) Selected images of (from top to bottom) dividing control, *ensc*, Klp67A-OE and Klp10A-OE NBs expressing tubulin (magenta) and myosin regulatory light chain (green and lower panels in monochrome) after anaphase onset (t=20s) till late telophase. Scale bar: 10μm. Time is s. B) Scheme showing the possible apical shift between the initial and final furrow curvature analysis. C) Dot plot showing of the relative furrow displacement between early anaphase and late telophase in control (0.00±0.04, *n=*18), *ensc* (−0.03±0.05, *n=*18), Klp67A-OE (−0.05±0.04, *n=*12), and Klp10A-OE NBs (−0.14±0.04, *n=*22). *: *P*<0.05, ***: *P*<0.001, ****: *P*<0.0001 (Wilcoxon test). D) Scheme showing the furrow width (red) during mid anaphase. E) Dot plot showing the relative myosin furrow width/cell length ratio for control (0.08±0.02, *n=*18), *ensc* (0.14±0.07, *n=*17), Klp67A-OE (0.13±0.05, n=12), and Klp10A-OE NBs (0.25±0.11, *n=*23), *: *P*<0.05, ***: *P*<0.001, ****: *P*<0.0001 (Wilcoxon test)).

### Centralspindlin is spatially and temporally regulated as two distinct populations

Our previous perturbations, which interfered with MT dynamics, suggested that a common mechanism was at play for maintaining the furrow position. In all higher eukaryotes examined to date, myosin recruitment and activation at the cleavage furrow is regulated by the highly conserved centralspindlin complex, a tetramer comprised of a Kinesin-6 family member complexed with Mgc-RacGAP (Pavarotti-klp and Tumbleweed in *Drosophila*, respectively)^7, 8, 47^. Strikingly, we found that the combination of Klp10A-OE and a single copy of the *pav*^*B200*^ null allele enhanced the asymmetry defect observed with Klp10A-OE alone (Figure S2 C). Centralspindlin functionality and targeting to the membranes is regulated by the chromosomal passenger complex (CPC)-dependent oligomerization ^48^. We therefore challenged the complex by introducing a single null allele for its Survivin subunit, *svn*^*2180*^, and monitored the effects on cell symmetry in the Klp10A-OE background. We found that Klp10A-OE-dependent size asymmetry defects were further enhanced when Survivin levels were reduced (Figure S2 C). These results suggest that the observed asymmetry defects are due, at least in part, to impaired centralspindlin function.

In most eukaryotic cells, the centralspindlin complex is located at the spindle midzone and at the equatorial cortex. To characterize the furrow mis-positioning that accompanies defective microtubule growth, we analyzed the spatio-temporal distribution of the motor component of centralspindlin, Pavarotti-klp, in different experimental backgrounds. We began by examining GFP-Pav-klp ^21^ localization in control NBs. Our time-lapse studies showed that most of the GFP-Pav-klp was located at the cortex at the cleavage site. Following the onset of furrow ingression, a second pool started to accumulate into a small and spatially distinct band near the former site occupied by the metaphase chromosomes at the spindle midzone (Figure 4 A, Figure S3A, and Video 7). The spatial and temporal separation of the GFP-Pav-klp signals led us to speculate that these were separate pools of centralspindlin. To confirm this hypothesis, we tracked GFP-Pav-klp in cells lacking MTs that were forced into anaphase using *Mad2* RNAi to abrogate the spindle assembly checkpoint ^14^. Under these conditions GFP-Pav-klp showed a slight enrichment at the basal cortex but this pool remained at almost baseline levels compared to control cells, which showed continuous recruitment of GFP-Pav-klp following anaphase onset (Figure S3 B and C). When microtubule polymerization was impaired in *ensc*, Klp10A-OE and Klp67A-OE cells, even if the centralspindlin component GFP-Pav-klp was initially present at the equatorial cell cortex, it did not become enriched at the cleavage site to the levels measured in controls (Figure 4 B, C, E red triangles, Figure S4 A, see also Video 8 and 9). Instead, in Klp10A-OE (Video 9), Klp67A-OE but not in *ensc* NBs (Video 8), GFP-Pav-klp appeared more abundant at the spindle midzone (Figure 4 B and C, see time 100 s blue arrows and insets at time 180 sec, Figure 4 E and Figure S4 B). Together, these experiments show that centralspindlin exists as two distinct and separable populations, one at the basal cortex and one at the spindle midzone. The decrease of the cortical centralspindlin pool is always accompanied by a displacement of the cleavage furrow.

**Figure 4.**
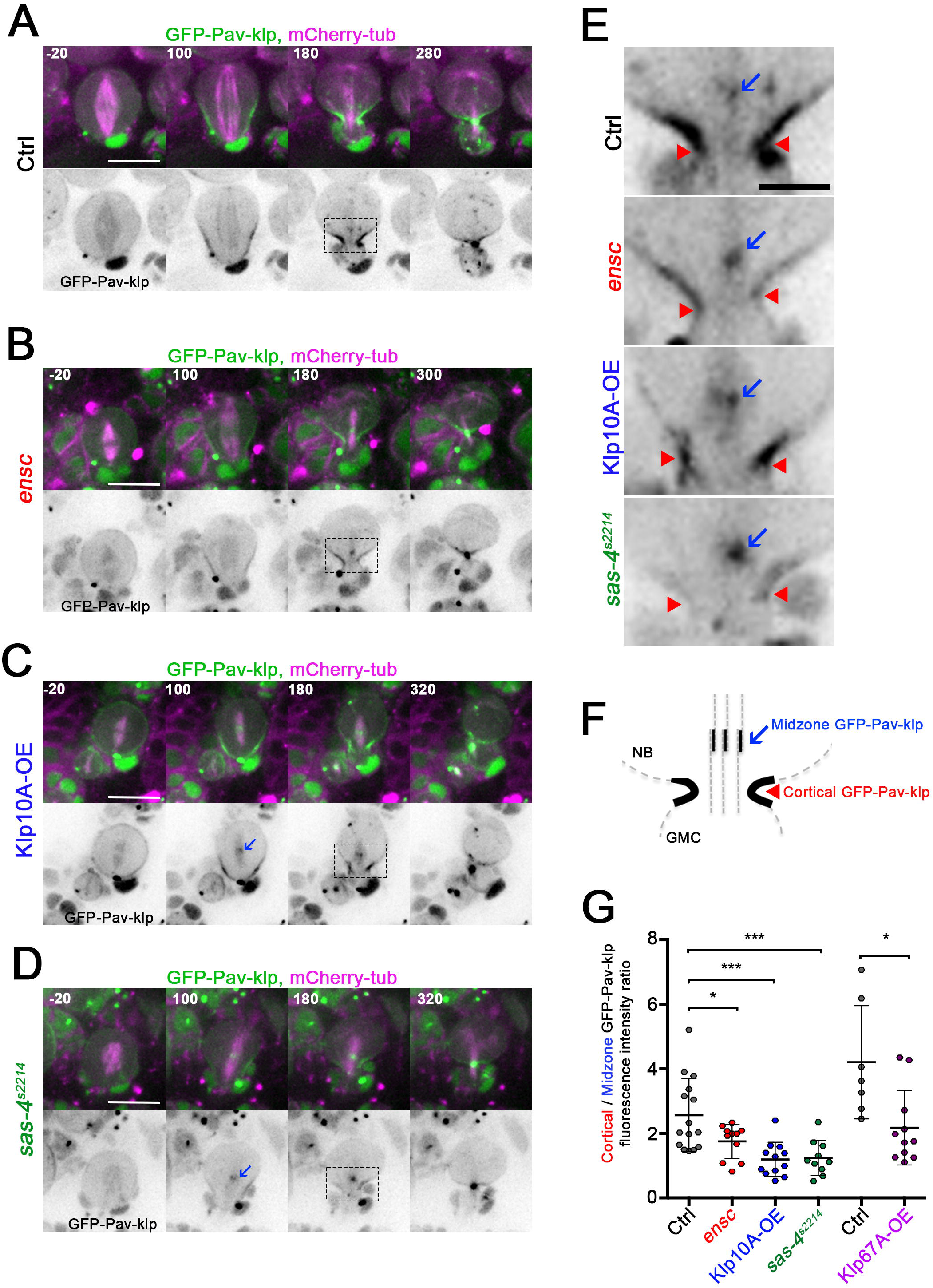
Analysis of centralspindlin localization and dynamics in control, *ensc*, Klp10A-OE and *sas-4*^*s2214*^ NBs. A) Selected images of dividing control expressing tubulin (magenta) and GFP-Pav-klp (green and lower panels in monochrome) from anaphase onset till late telophase (top left). B) *ensc* NB. C) Klp10A-OE NB. D) *sas-4*^*s2214*^ NBs (bottom). Scale bar: 10μm. Time is s. E) Higher magnification view of the selected control *ensc*, Klp10A-OE and *sas-4*^*s2214*^ telophase NBs (from panels in A-D) showing GFP-Pav-klp localization at the cleavage site. See the strong signal at the cell cortex (red arrowheads) and the weak signal at the presumptive spindle midzone away from the cleavage site, toward the apical side blue (blue arrows). F) Scheme of the cleavage site showing the cortical and midzone centralspindlin pools. G) Dot plot (± s.d.) showing the relative cortical/midzone GFP intensity ratio for control (2.57±1.13, *n=*15), *ensc* (1.75±0.52, *n=*11), Klp10A-OE (1.20±0.53, *n=*12), *sas-4*^*s2214*^ (1.24±0.54, *n=*10), control (4.2±1.76, *n=*7) and Klp67A-OE NBs (2.17±1.15, *n=*11). *: *P*<0.05, ***: *P*<0.001, ****: *P*<0.0001 (Wilcoxon test).

### The spatio-temporal regulation of centralspindlin relies on stable peripheral MTs

Fluorescence quantification (Figure 4 F) revealed that compared to wild type, Klp10-OE and Klp67A-OE NBs both displayed a decrease in cortical GFP-Pav-klp signal with a concomitant increase at the midzone (Figure S4). To further characterize the relationship between cortical and midzone centralspindlin pools and the role of MT growth in asymmetrical cleavage, we examined GFP-Pav-klp dynamics in *sas4*^*s2214*^ mutants, which lack centrosomes and their associated astral MTs ^28^. In this background, cortical enrichment also appeared diminished relative to the midzone (Figure 4 D, Figure S4 A and B). An enlarged view of the boxed regions for 180 sec post-anaphase onset highlights this increased centralspindlin recruitment at the spindle midzone and weaker accumulation at the cell cortex (Figure 4 E). Although *sas4*^*s2214*^ NBs exhibited signal enrichment at their midzones, not all cells had a clear cortical reduction (Figure S4). Strikingly, *sas4*^*s2214*^ mutants exhibited both an increased spindle length as well as a significant overall cell size asymmetry defect (Figure S5 A and B), further supporting the idea that astral MTs maintain basal furrow position. To confirm the contribution of the MT-aster in furrow positioning and maintenance, we removed it by laser ablation of the basal centrosome. Each centrosome was labeled with GFP-tagged Aurora A and basal proximal centrosome was ablated by a multi-photon laser until the signal was no longer detectable. Consistent with centrosome removal, ablated cells displayed a phenotype virtually identical to *sas4*^*s2214*^ mutants: daughter cells exhibiting cell size asymmetry after the ensuing cytokinesis (Figure S5 C and D, Video 10, compare left and right). These live cell observations suggested that astral MTs were essential to furrow positioning. We therefore performed a quantitative analysis of fixed preparations examining MT distribution during early telophase. Detailed morphological examination revealed that in wild type NBs bundles of astral MTs capped the future ganglion mother cell and spread apically, closely apposed to the cortex at the cleavage furrow. This was not the case with *ensc*, Klp67A-OE or Klp10-OE NBs, which showed decreased MT densities and lacked the presumptive bundles (Figure S5 E and F). Altogether, our data strongly suggest that peripheral astral MTs originating from the basal centrosome, in a close vicinity of the basal furrow play a key role in accurate asymmetric cell division.

### The early spindle midzone and furrow occupy distinct positions in NBs

The presence of GFP-Pav-klp at the spindle midzone distal to the cleavage site and the movement of the furrow towards the equator in peripheral MTs-deficient cells prompted us to further characterize the cleavage site and the midzone in wild type cells. For this purpose, we used Fascetto-GFP (the homologue of the mammalian PRC1 protein; Feo-GFP) a marker that uniquely labels the spindle midzone (Figure 5 A) ^27, 49^. We found that in these NBs the metaphase plate was slightly shifted toward the basal side relative to the cell equator along the apico-basal axis (Figure 5 A, −120 sec, arrowhead and Figure 5 C) but similarly placed to the midzone-defining Feo-GFP signal that appears following anaphase chromosome segregation (Figure 5 A, time 90 sec, and Figure 5 D arrowhead). This was in contrast to the position of the furrow (Figure 5 A, time 90, compare green and white arrows; E arrowhead). Indeed, while the metaphase plate and midzone were interchangeably located, the furrow was always distinct and basally distal to these (Figure 5 F, Video 11). Interestingly, kymograph analyses of the spindle midzone and cell membranes reveals that the midzone moves basally during the ingression of the furrow until they ultimately consolidate into a single structure (Figure 5 B). In summary, these data demonstrate that the spindle midzone in wild type cells occupies a spatially different position than that of the furrow and its associated cortical MTs. Both of these can recruit centralspindlin, however, under normal circumstances it is the cortical pool that dominates in *Drosophila* NBs to define the cleavage site.

**Figure 5.**
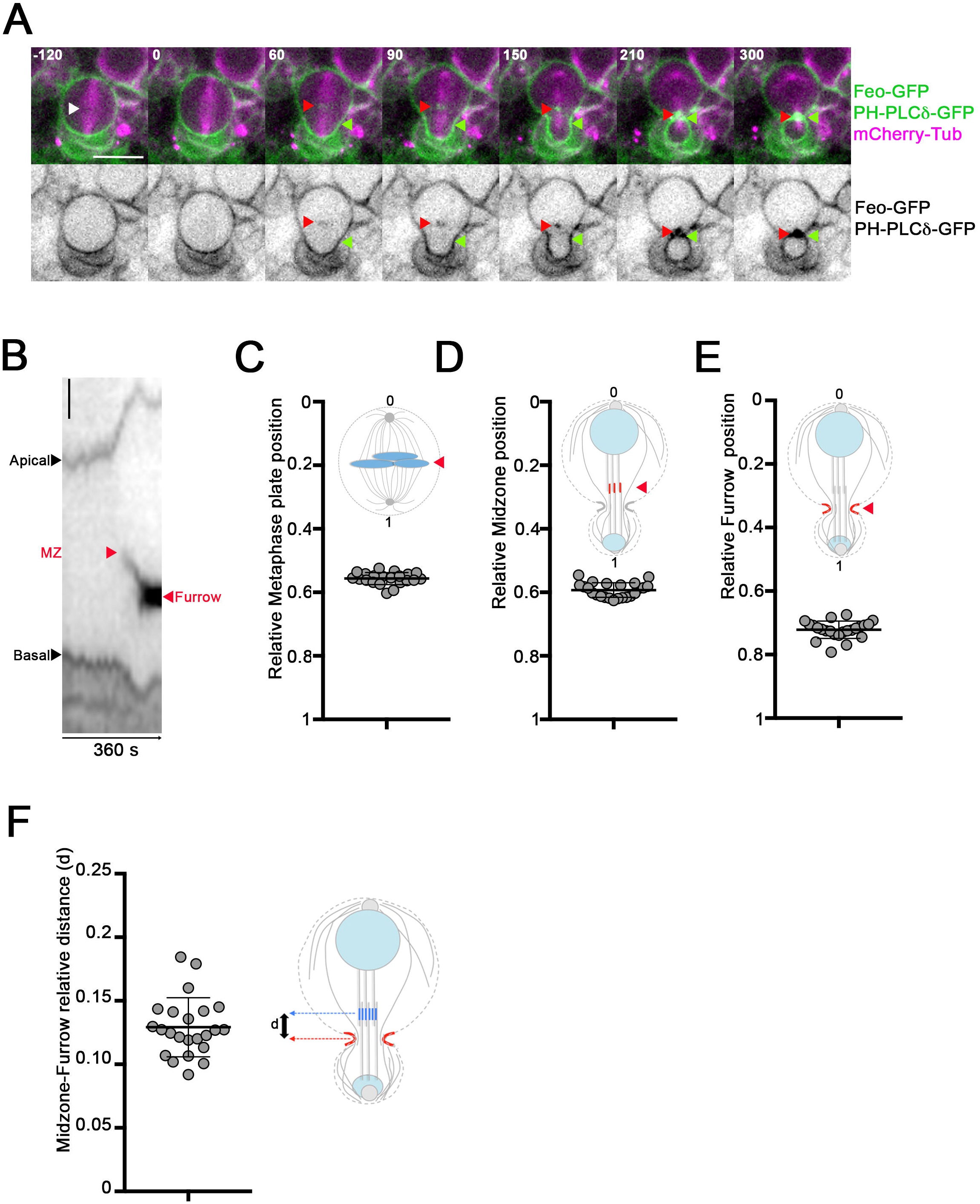
Analysis of the spindle midzone and the cleavage furrow position in brain NBs during cell division. A) Selected images of a WT NB expressing Feo-GFP (green and lower panels in monochrome), MTs (magenta), and membranes during cell division (green and lower panels in monochrome). The metaphase plate is indicated by a white arrowhead (−120s). The spindle midzone is indicated by red arrowheads and the furrow is indicated by green arrowheads (60-300s). Scale bar: 10 μm. Time is s. B) Kymograph showing the localization of the spindle midzone and the cell contours during the time course of the NB cell division shown in panel A, along the apico-basal axis. C) Dot Plot (± s.d.) showing the mean relative metaphase plate position along the apico-basal cortex 90 sec after anaphase onset (0.56± 0.02 *n*=21). D) Dot Plot showing the mean relative spindle midzone position (± s.d.) along the apico-basal cortex 90 sec after anaphase onset (0.59±0.02, *n*=23). E) Dot Plot (± s.d.) showing the mean relative furrow position along the apico-basal cortex 90 sec after anaphase onset (0.72±0.13, *n*=23). F) Dot Plot (± s.d.) showing the mean relative distance between the furrow position and the spindle midzone 90 sec after anaphase onset (0.13 ± 0.02, *n*=23).

## Discussion

Asymmetric cell division is a robust process that ensures that two daughter cells inherit different fates and sizes. The *Drosophila* NB is a powerful and widely used model system to study this specialized form of division because of the large number of NBs in the developing *Drosophila* brain, rapid division time and experimental traceability ^50-52^. Moreover, although these cells are relatively small, they are highly asymmetrical following cytokinesis allowing accurate measurements and analyses. In this study, we have used this model system to challenge asymmetric cell division after modification of MT growth dynamics. We were able to increase mitotic spindle length using over-expression of MT polymerizing MAPs (Msps and Ensconsin), as well as by RNAi-mediated depletion of Klp67A, a member belonging to Kinesin-8 family of MT depolymerizing Kinesins. Despite the presence of long and bent mitotic spindles under these conditions, the NB cell size ratio remained unchanged relative to wild type NBs. This reveals that asymmetric cell division and asymmetric positioning of the cleavage furrow are resistant to an excess of abnormally long and stable MTs during cell division. By contrast, decreasing MT stability and shortening of the mitotic spindle produced more symmetric cell divisions. This change was due to an apical shift of the cleavage furrow during its ingression. This phenotype was not MAP specific since over-expression of either Klp10A (Kinesin-13) or Klp67A (Kinesin-8) MT depolymerases, as well as the deletion of *Ensconsin* produced similar effects. Rather they suggest that spindle size or interference with microtubule dynamics is responsible for the phenotype. Interestingly, *sas-4*^*s2214*^ mutants which are reported to lack functional centrosomes and thus astral microtubules yielded reduced levels of cell size asymmetry although these NBs harbored longer mitotic spindles ^28^. This reveals that astral MTs and not the mitotic spindle length is the key element responsible for the level of size asymmetry observed in NBs. Consistent with this hypothesis, loss of the basal MT-aster, through targeted laser irradiation and ablation prior anaphase onset, also reduced sibling cell size asymmetry. Taken together these results strongly suggest that a population of basal peripheral astral MTs is required to maintain a cleavage site, which normally favors a basal position in the fly neuroblast. In agreement with this our quantification of peripheral MT bundles close to the ingression furrow revealed that they are significantly decreased during telophase in *ensc*, Klp10A-OE and Klp67A-OE NBs. Our results are in accord with reports indicating that a subpopulation of stable astral MTs play a key role in the initiation of furrowing in symmetrically dividing cells and that in some systems, furrowing can occur without the presence of a stable central spindle ^3, 6, 7, 20, 53-56^.

However in contrast to other studies which utilize micromanipulation and laser ablation, our data reveals that in wild type asymmetrically dividing NBs, the astral MT furrowing pathway dominates over the midzone pathway. Previous studies have suggested that NBs have two genetically separable pathways to drive cytokinesis. The first, cortical polarity pathway, is responsible for the targeting of the furrowing machinery to the basal cortex on the surface of what is destined to become the smaller differentiating ganglion mother cell ^11, 12^. Interference with this polarity pathway prevents myosin enrichment at the basal cortex, leading to symmetric division. The alternative spindle pathway relies on the spindle midzone and the chromosomal passenger complex ^9^,^12^. However, several of our observations are at odds with these previous findings and favor a model of furrow positioning that relies largely on peripheral astral MTs with little if any contribution from the spindle midzone; (i) live cell imaging and analyses utilizing GFP-Pav-klp as a marker of centralspindlin position revealed that this master controller of cytokinesis accumulated at the basal cortex throughout the entire furrow ingression process. (ii) Centralspindlin levels were low at the midzone during furrow placement and ingression compared to the cortex (Figure 4). (iii) We consistently observed that the midzone, as defined independently using both GFP-Pav-klp and Feo-GFP, was spatially independent from the furrowing site (Figures 5B, 6A). In addition, the midzone ultimately moved to the position of the furrow and not vice-versa (Figure 5B), confirming previous observations in embryonic NBs ^57^. (iv) Finally, genetic or photo-based removal of the basal centrosome precluded peripheral MT formation and interaction with the cortex. Accordingly, the cortical centralspindlin pool was diminished and cells experienced a size asymmetry defect (Figure 4 and S4).

Our localization studies employing Feo-GFP suggest that under normal conditions midzone-associated centralspindlin does not perform a key role in positioning of the cleavage site and that this function is served by the more abundant centralspindlin pool associated with the membranes at the cleavage site. However, when peripheral MTs were impaired, centralspindlin enrichment at the furrow was diminished, leading to a decreased midzone/furrow centralspindlin ratio and a reset of the furrowing toward the equatorial midzone. This indicates that the two populations of centralspindlin are competent to signal furrowing but that the cortical pool delivered by astral MTs is normally dominant. Thus the spatial localization and the cortical/midzone ratio of centralspindlin are the pivotal determinants of final furrow positioning in the *Drosophila* NB. Interestingly, a recent study has shown that a similar competition between centralspindlin pools also occurs in human cells ^58^, revealing an evolutionary conservation of the mechanism.

As with human cells, we found that the CPC activity seems essential in this regulatory event (Figure S2 C). In contrast to a recent study in S2 cells, we do not observe GFP-Pav-klp labelling at the plus ends of astral MTs ^59^ even when studied by enhanced resolution imaging methods. Instead, we consistently find that centralspindlin coats the entire length of astral MTs, suggesting that the plus end directed motor activity of Pav-klp is used to bring centralspindlin to the furrow in *Drosophil*a NBs in agreement with previous studies in early embryos ^21, 60^.

Altogether, our data suggest a model in which a competition between different centralspindlin populations is a key determinant of asymmetric division in *Drosophila* NBs. The concerted and consecutive action of polarity cues and astral MTs as centralspindlin delivery arrays are essential in this process. The ability of the spindle midzone to define furrow and cleavage location only becomes engaged during late telophase and after subcortical astral MTs are compromised. Despite their clear role in governing size asymmetry, we have not been able to induce complete daughter cell size equality through any of a host of MT perturbing treatments. This suggests that additional feedback mechanisms exist to ensure a threshold level of asymmetry in these cells. Elucidating these systems and their evolutionary advantages will be important directions for future investigations.

## Acknowledgments

We thank Gregory Rogers, Pier Paolo d’Avino, Renata Basto, Gohta Goshima, Jordan Raff, Hiro Ohkura, Anne Royou, Roger Karess, Christian Dahmann, Antoine Guichet, Juliette Mathieu, Jean-René Huynh, Clemens Cabernard, Tri Pham for providing fly stocks, antibodies and cDNAs. We thank Chloé Rauzier for preliminary functional analyzes of Ensc-OE and Msps-OE NBs. This work was funded by the Ligue Nationale Contre le Cancer, the Fondation ARC pour la Recherche sur le Cancer. A. T. is a doctoral fellow of the Région Bretagne and the Ligue Nationale contre le Cancer. We thank the Photonic Imaging Center of Grenoble Institute of Neurosciences, which is part of ISdV core facility. We thank Xavier Pinson, Stéphanie Dutertre and Sébastien Huet for advices and help with the microscopes and the Microscopy Rennes Imaging Center platform. We thank Romain Gibeaux, Pier Paolo d’Avino and Christelle Benaud for ideas, critical readings and helpful suggestions. The authors have no competing financial interests to declare.

---

**Figure S1.**
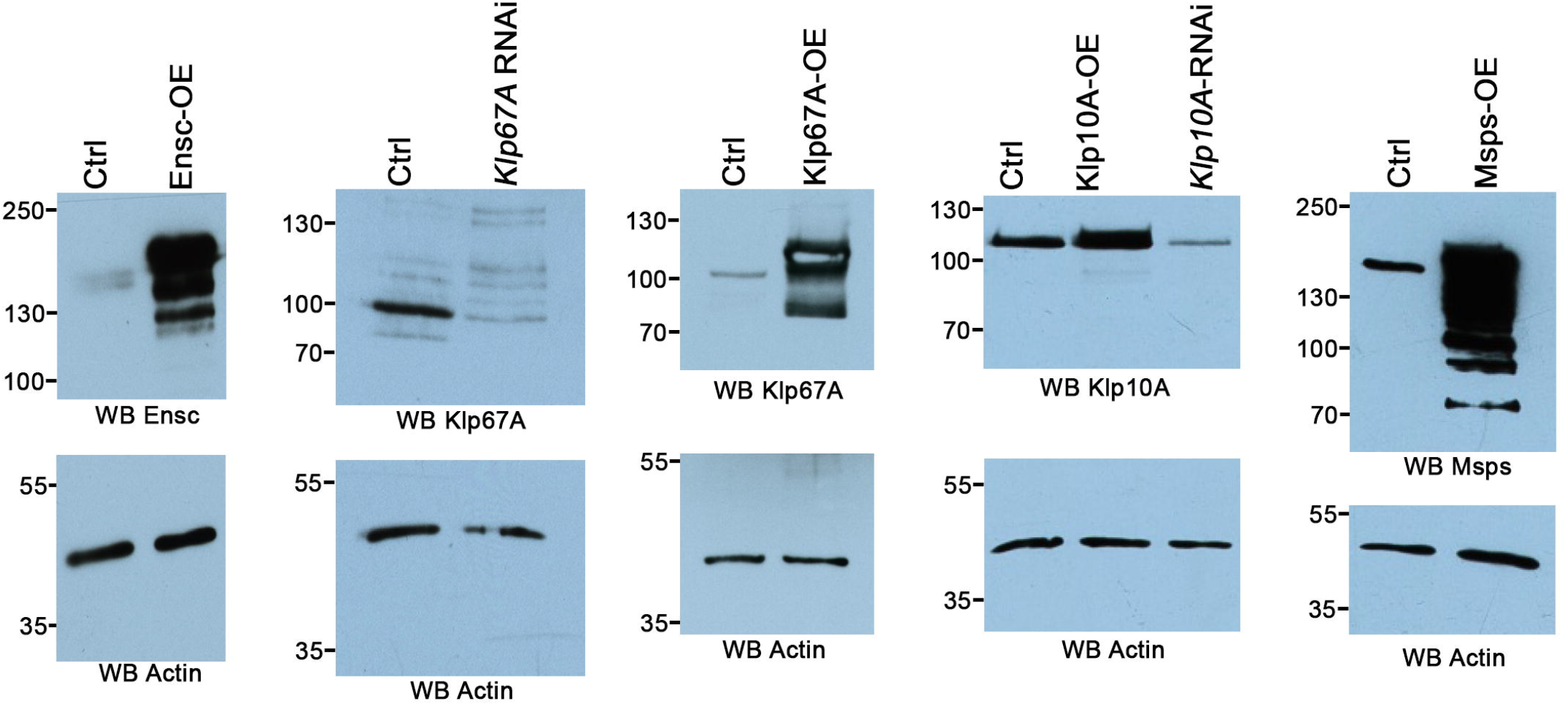
Western Blot analyses of Ensconsin, Minispindles, Klp10A, Klp67A protein in brain extracts. Brain subjected to RNAi or overexpression (OE) for the indicated MAPs were dissected and resuspended in sample buffer before for Western blot analyses. The primary antibody used for each Western blot is indicated at the bottom of each top panel. In the bottom panels, actin was used as a loading control.

**Figure S2.**
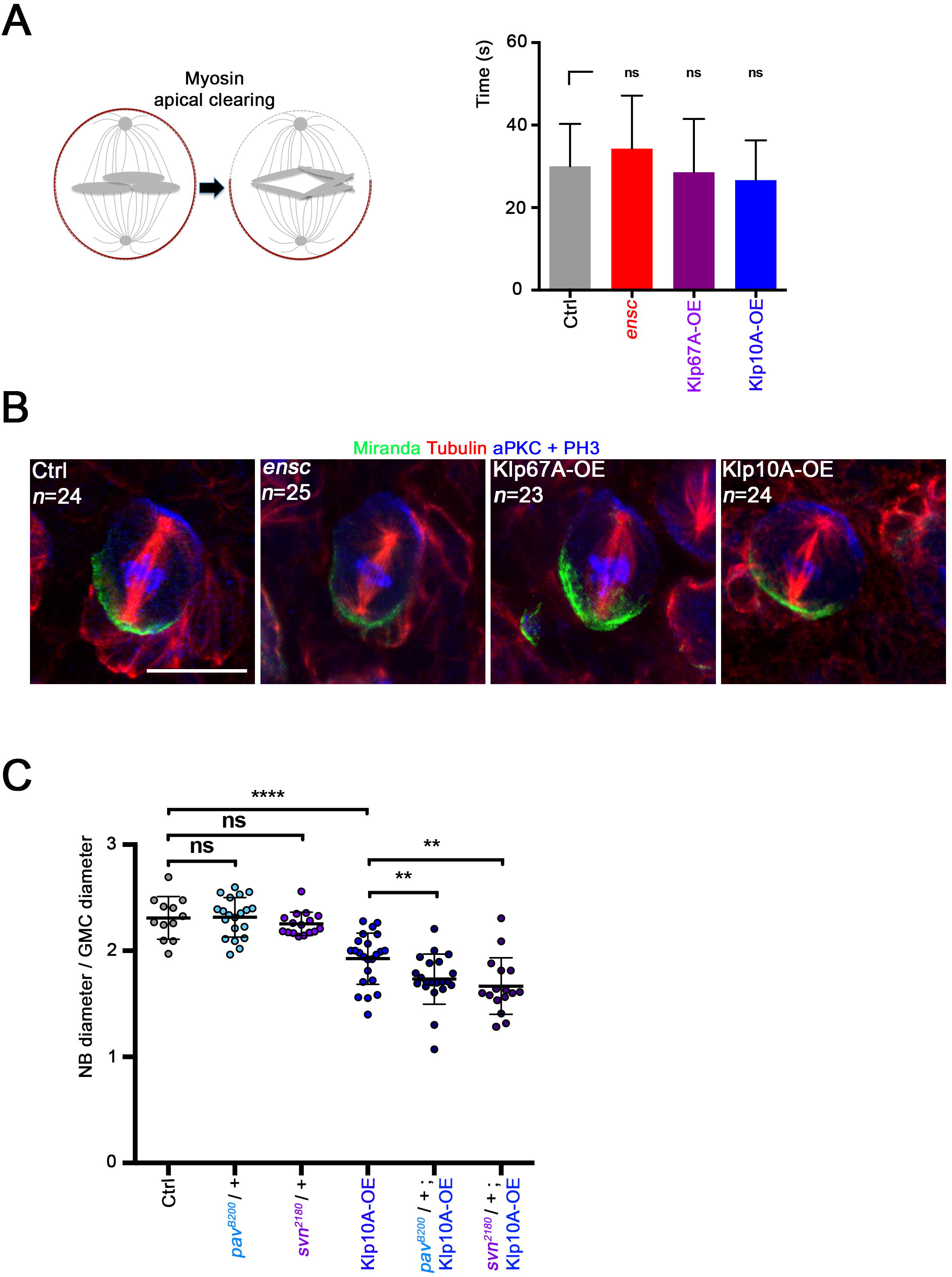
Polarity-dependent apical myosin clearing and localization of aPKC and Miranda in *ensc*, Klp10A-OE and Klp67A-OE NBs. A) Scheme of Myosin clearing from the apical cortex of neuroblast at metaphase/anaphase transition (left). Histogram showing the timing of apical myosin clearing (right) for NBs of the indicated genotypes in control (30.00±10.29 s, *n=*18), *ensc* (34.29±12.87 s, *n=*21), Klp67A-OE (28.57±12.92 s, *n=*14), Klp10A-OE (26.67±9.63 s, *n=*24) NBs. Time 0s is anaphase onset. ns: non-significant. (Wilcoxon test) B) Analysis of aPKC and Miranda localizations in control, Klp10A-OE, Klp67A-OE and in *ensc* metaphase NBs. Brains of the indicated genotypes were fixed and stained for aPKC (blue), Miranda (green), α-tubulin (red) and phospho-histone H3 (Blue). Scale bar: 10μm. The number of examined cells (n) is also indicated. None of the conditions used compromise the location of aPKC and Miranda. C) Dot plot showing the NB diameter/GMC diameter ratio (± s.d.) in control (2.31±0.20, *n=*12), in *pav*^*B200*^/+ (2.32±0.19, *n*=20), in *svn*^*2180*^/+ (2.25±-0.11, *n=*16), in Klp10A-OE (1.93±0.24, *n=*23), in *pav*^*B200*^/+; Klp10A-OE (1.73±0.24, *n=*21), and in *svn*^*2180*^/+; Klp10A-OE (1.67±0.27, *n=*16) NBs. ns: non-significant, **: *P*<0.01, ****: *P*<0.0001 (Wilcoxon test). Cell size asymmetry in Klp10A-OE NBs is enhanced by alteration of the CPC and the centralspindlin complex.

**Figure S3.**
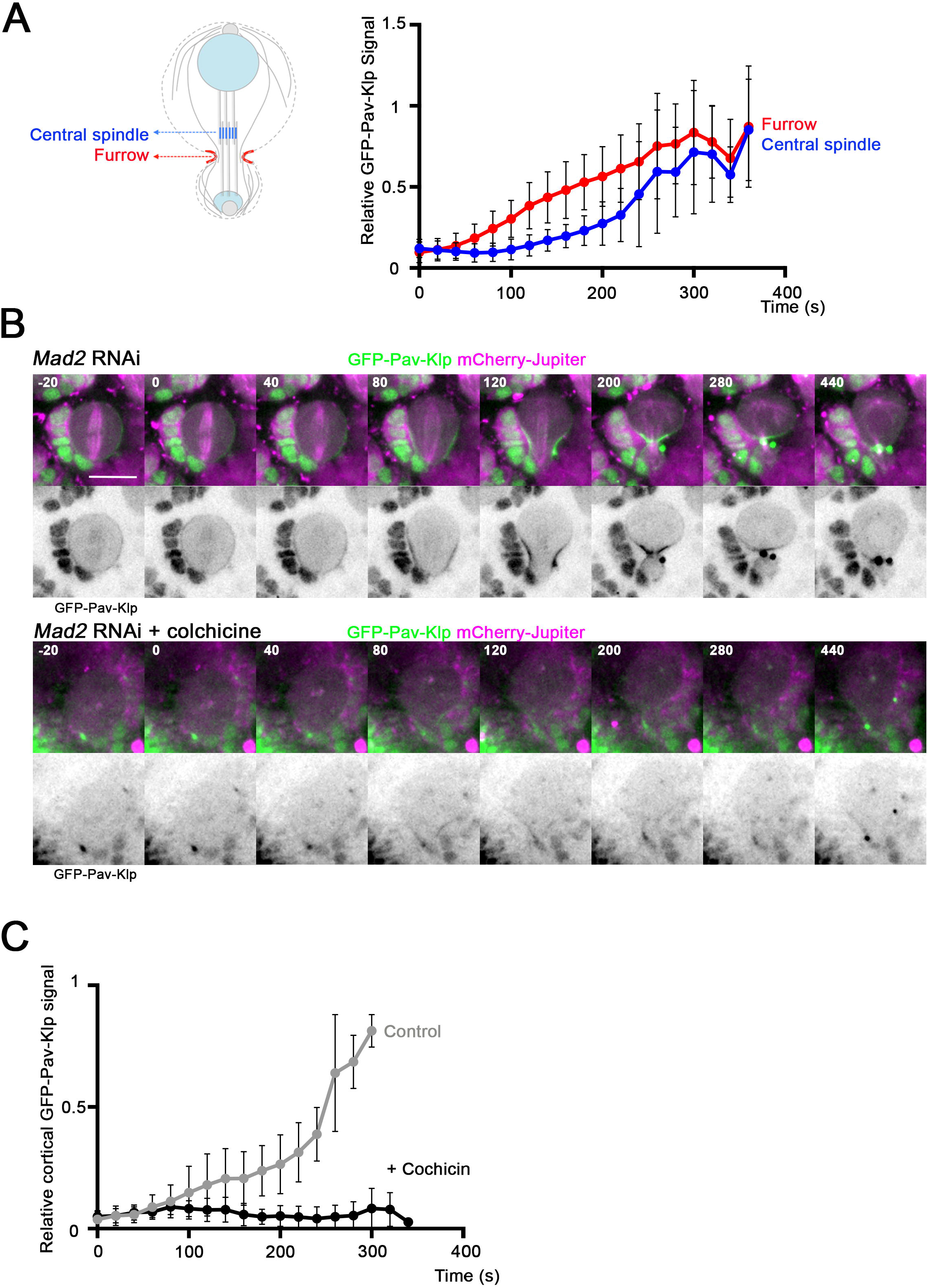
Midzone and cortical centralspindlin recruitment in control and colchicine-treated NBs. A) Mean relative cortical/furrow (red) and central spindle (blue) enrichment of Pav-klp-GFP (± s.d.) in a control NB (regarding Figure 4 A). Pav-klp-GFP is rapidly recruited at the cortex while its recruitment at the centralspindle is slower. *n*=15. B) GFP-Pav-klp (green and lower panels in monochrome) is strongly recruited to the cell cortex and to the cleavage furrow in Mad2-depleted cells. In the absence of MTs (*Mad2* RNAi+colchicine), GFP-Pav-klp is faintly detected at the basal cortex but is not enriched similarly to conditions where the MT cytoskeleton n is intact. Scale bar: 10μm. Time is s. C) Mean relative GFP-pav-Klp signal (± s.d.) at the cell cortex in *Mad2* RNAi with (*n*=9) (grey) or without (*n*=7) (dark grey) MTs.

**Figure S4.**
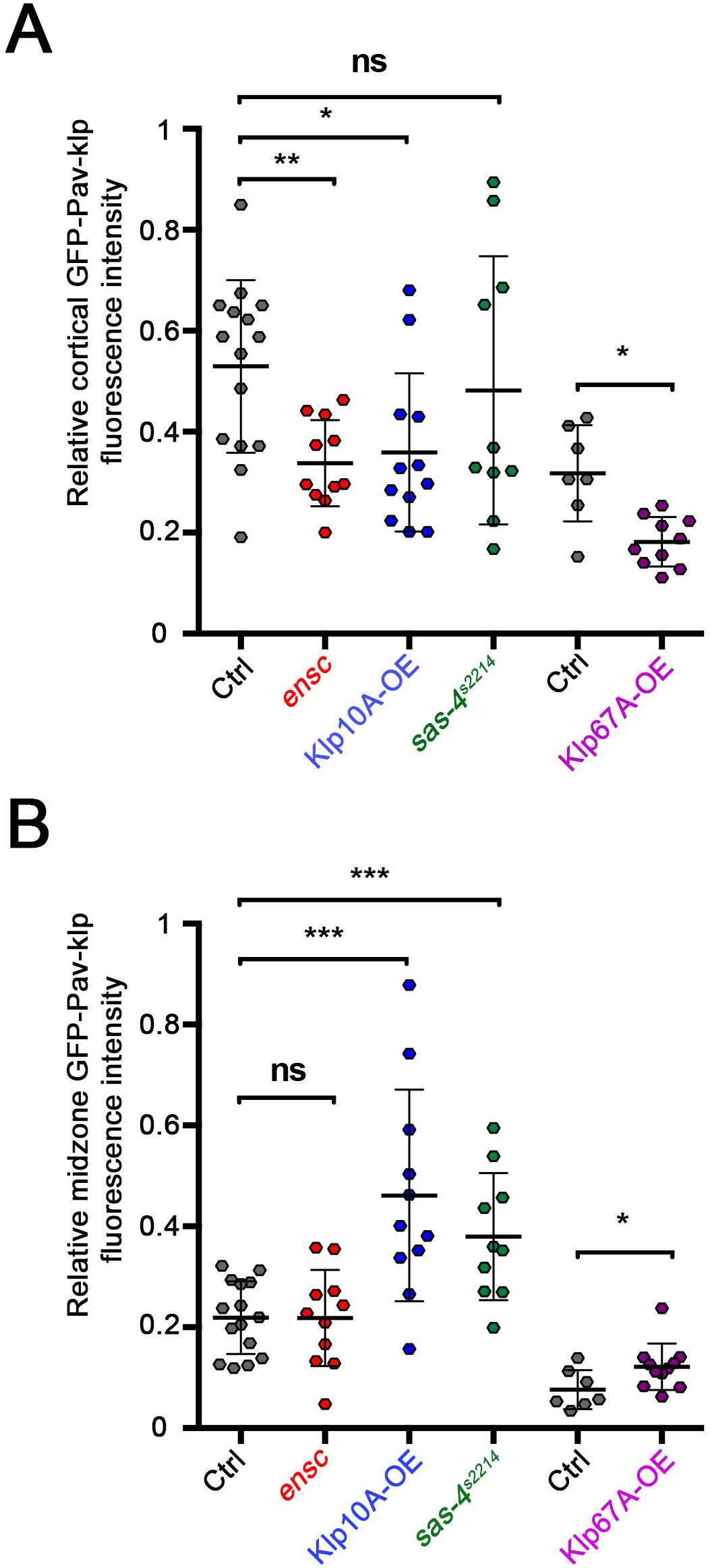
Analysis of centralspindlin levels at the centralspindle or the cleavage furrow during anaphase. A) Dot plot showing the mean relative GFP-Pav-klp signal at the cleavage furrow 180 sec after anaphase onset in control (0.53±0.17, *n=*15), *ensc* (0.34±0.09, *n=*11) Klp10A-OE (0.36±0.16, *n=12*), and *sas4*^*s2214*^ (0.48±0.27, *n=*10), control (0.32±0.10, *n=*7), and Klp67A-OE (0.18±0.05, *n=*11) NBs. ns: non-significant, ****: *P*<0.0001, (Student test and Wilcoxon test fort Klp67A-OE). B) Dot plot showing the total relative GFP-Pav-klp signal at the midzone 180 sec after anaphase onset in control (0.22±0.07, *n=*15), *ensc* (0.224±0.09, *n=*11) Klp10A-OE (0.46±0.20, *n=12*), and *sas4*^*s2214*^ (0.38±0.13, *n=*10), control (0.08±0.04, *n=*7), and Klp67A-OE (0.12±0.05, *n=*11) NBs. ns: non-significant, *: *P*<0.05, **:*P*<0.01, ****: *P*<0.0001, (Student test and Wilcoxon test for Klp67A-OE).

**Figure S5.**
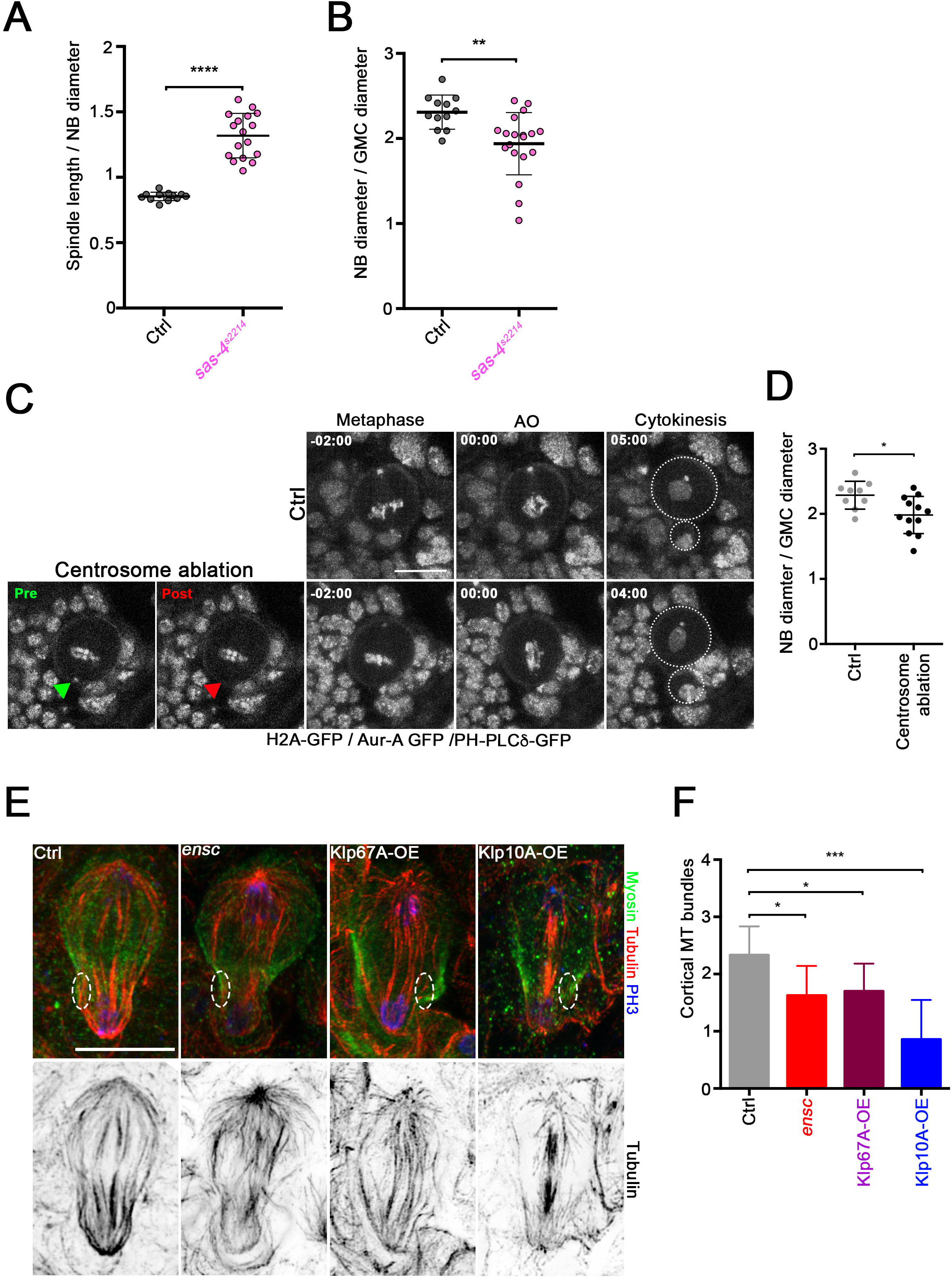
Analysis of centrosome contribution for cleavage positioning and of peripheral MTs in different backgrounds. A) Dot plot showing the mitotic spindle length/NB diameter ratio (± s.d.) in control (0.85±0.03, *n=*12) or *sas4*^*s2214*^ NBs (1.32±0.17, *n=*17), ****: *P*<0.0001, (Wilcoxon test) B) Dot plot showing the NB diameter/GMC diameter ratio (± s.d.) in control (2.31±0.2, *n=*12) or *sas4*^*s2214*^ NBs (1.94±0.37, *n=*19). **: *P*<0.01 (Wilcoxon test). C) Selected images of control NBs (top) or NBs in which the basal centrosome has been laser-ablated (bottom). The green triangle shows the basal centrosome before ablation, the red triangle shows the position of the ablated centrosome. The NBs expressed GFP-H2A (to confirm the metaphase stage, anaphase onset and the position of the basal GMC), PH-PLCδ-GFP (to visualize cell contours) and GFP-Aurora A (to target the centrosomes). D) Dot plot showing the NB diameter/GMC diameter ratio (± s.d.) in control (2.29±0.21, *n=9*) or basal centrosome-ablated NBs (1.98±0.29, *n=*12). ***: *P*<0.001, (Student test). Scale bar: 10μm. Time is min:s. E) Selected high-resolution images of control, *ensc*, Klp67A-OE, and Klp10A-OE NBs. Tubulin is shown in red (and lower panels in monochrome), Myosin is shown in green, and phospho-Histone H3 (Ser10) in blue. The white ellipses show the peripheral MT bundles in the vicinity of the furrow. Scale bar: 10μm. F) Histogram showing the number of peripheral astral MTs bundles detected at the vicinity of the middle cleavage plane in a Z series projection (1 μm) in control (2.3±0.5, *n=*9), *ensc* (1.63±0.52, *n=*8), Klp67A-OE (1.7±0.48, *n=*10) and in Klp10A-OE (0.86±0.69, *n=*7) NBs. *: *P*<0.05, ***: *P*<0.001 (Wilcoxon test).

**Video 1. Asymmetric cell division in a control NB**

Microtubules (RFP-α-tubulin) are displayed in magenta and PH-PLCδ-GFP is displayed in green. Scale bar: 10 μm. Time is min: s.

**Video 2. Asymmetric cell division in a Msps-OE NB**

Microtubules (Msps-RFP) are displayed in magenta and PH-PLCδ-GFP is displayed in green. Scale bar: 10 μm. Time is min: s.

**Video 3. Asymmetric cell division in a Klp10A-OE NB**

Microtubules (RFP-α-tubulin) are displayed in red and PH-PLCδ-GFP is displayed in green. Scale bar: 10 μm. Time is min: s.

**Video 4. Cortical myosin dynamics during cytokinesis in a control NB**

Microtubules (mCherry-α-tubulin) are displayed in magenta and Sqh-GFP is displayed in green or monochrome (right). Scale bar: 10 μm. Time is min: s.

**Video 5. Cortical myosin dynamics during cytokinesis in an ensc NB**

Microtubules (mCherry-α-tubulin) are displayed in magenta and Sqh-GFP is displayed in green or monochrome (right). Scale bar: 10 μm. Time is min: s.

**Video 6. Cortical myosin dynamics during cytokinesis in a Klp10A-OE NB**

Microtubules (mCherry-α-tubulin) are displayed in magenta and Sqh-GFP is displayed in green or monochrome (right). Scale bar: 10 μm. Time is min: s.

**Video 7. Pavarotti-klp localization during cytokinesis in a control NB**

Microtubules (mCherry-α-tubulin) are displayed in magenta and GFP-Pav-klp is displayed in green or monochrome (right). Scale bar: 10 μm. Time is min: s.

**Video 8. Pavarotti-klp localization during cytokinesis in an ensc NB**

Microtubules (mCherry-α-tubulin) are displayed in magenta and GFP-Pav-klp is displayed in green or monochrome (right). Scale bar: 10 μm. Time is min: s.

**Video 9. Pavarotti-klp localization during cytokinesis in a Klp10A-OE NB**

Microtubules (mCherry-α-tubulin) are displayed in magenta and GFP-Pav-klp is displayed in green or monochrome (right). Scale bar: 10 μm. Time is min: s.

**Video 10. Asymmetric cell division in a control NB or after basal centrosome ablation**

The left panel shows a control NB. The right panel shows a NB in which the basal centrosome was ablated. First and second images show pre and post ablation images respectively before acquisition. Aurora A-GFP, GFP-H2A and PH-PLCδ-GFP are displayed in grey. Scale bar: 10 μm. Time is min: s.

**Video 11. Fascetto localization during cytokinesis in a control NB**

Microtubules (mCherry-α-tubulin) are displayed in magenta and PH-PLCδ-GFP and Feo-GFP are displayed in green or monochrome (right). Scale bar: 10 μm. Time is min: s.

